# Missing Wedge Completion via Unsupervised Learning with Coordinate Networks

**DOI:** 10.1101/2024.04.12.589090

**Authors:** Dave Van Veen, Jesús G. Galaz-Montoya, Liyue Shen, Philip Baldwin, Akshay S. Chaudhari, Dmitry Lyumkis, Michael F. Schmid, Wah Chiu, John Pauly

**Affiliations:** Dept. of Electrical Engineering, Stanford University; Dept. of Bioengineering, Stanford University; Dept. of Electrical and Computer Engineering, University of Michigan; Dept. of Biochemistry and Molecular Pharmacology, Baylor College of Medicine; Dept. of Genetics, The Salk Institute for Biological Sciences; Dept. of Radiology, Stanford University; Graduate School of Biological Sciences, University of California San Diego; Division of CryoEM and Bioimaging, SSRL, SLAC National Accelerator Laboratory; Dept. of Microbiology and Immunology, Stanford University

## Abstract

Cryogenic electron tomography (cryoET) is a powerful tool in structural biology, enabling detailed 3D imaging of biological specimens at a resolution of nanometers. Despite its potential, cryoET faces challenges such as the missing wedge problem, which limits reconstruction quality due to incomplete data collection angles. Recently, supervised deep learning methods leveraging convolutional neural networks (CNNs) have considerably addressed this issue; however, their pretraining requirements render them susceptible to inaccuracies and artifacts, particularly when representative training data is scarce. To overcome these limitations, we introduce a proof-of-concept unsupervised learning approach using coordinate networks (CNs) that optimizes network weights directly against input projections. This eliminates the need for pretraining, reducing reconstruction runtime by 3 − 20× compared to supervised methods. Our *in silico* results show improved shape completion and reduction of missing wedge artifacts, assessed through several voxel-based image quality metrics in real space and a novel directional Fourier Shell Correlation (FSC) metric. Our study illuminates benefits and considerations of both supervised and unsupervised approaches, guiding the development of improved reconstruction strategies.

## 1 Introduction

Recent advancements in cryogenic electron microscopy (cryoEM) [1, 2] have elevated it from a specialized technique to a cornerstone of structural biology [3, 4] and molecular sciences [5]. Cryogenic electron tomography (cryoET), an extension of cryoEM, offers detailed three-dimensional (3D) representations of macromolecules, cells, and tissues in states close to their natural environment at nanometer-scale resolution [6]. This technique is versatile, allowing for the examination of a wide array of macromolecular complexes *in vitro* [7] and *in situ* [8], including amyloid filaments [9–14] and enveloped viruses such as SARS-CoV-2 [15]. CryoET can also probe the structure of other clinically-relevant samples ranging from individual organelles [16, 17] and cells [18, 19] to complex tissue sections [20–24]. Insights from cryoET data analysis can facilitate understanding dynamic molecular processes such as viral infections [25–30], enable pathology diagnosis [31, 32], and reveal the impacts of potential therapeutic interventions [19], among other biomedical applications. A critical downstream technique, subtomogram averaging (STA) [33, 34], further refines cryoET data to achieve subnanometer [35] to near-atomic [36] resolution of repeated structures within tomograms [37, 38], underscoring the importance of precise cryoET reconstructions for accurate particle localization and structural analysis.

CryoET involves the rapid freezing of biological specimens followed by their examination with a transmission electron microscope (TEM) [39]. This process entails capturing a series of two-dimensional (2D) projection images as the specimen is incrementally tilted, compiling what is known as a tilt series [40]. These images are then aligned and combined to produce a 3D reconstruction, or tomogram, typically through weighted-back projection (WBP) methods [41], as enabled by software such as IMOD [42]. Despite cryoET’s ability to capture intricate structural details, the technique is limited by the specimens’ susceptibility to radiation damage [43] and the inherent mechanical constraints of TEM, which restrict the tilt series to angles between -60° and +60°. This limitation results in the “missing wedge” phenomenon [44], where the lack of data at experimentally inaccessible angles leads to artifacts that distort tomogram quality. This affects the resolution and density accuracy of visualized features, hence complicating 3D analyses. Such distortions are particularly problematic for structures perpendicular to the electron beam, often resulting in the omission of critical top and bottom details in images of spherical, oblong, or elongated biological features (*e*.*g*. cell membranes, organelles, vesicles, microtubules, and virions).

Efforts to mitigate the missing wedge’s impact have ranged from introducing new data collection techniques, such as dual-axis [45] and conical [46] tomography, to novel applications of statistical and iterative data processing methods [47–49], including total variation minimization [50] and compressed sensing [51]. In the realm of supervised deep learning, IsoNet stands out by employing a convolutional neural network (CNN) U-Net [52] trained on subtomograms extracted from tomograms reconstructed via weighted-back projection (WBP), intentionally adding missing wedge artifacts to create a paired training set. IsoNet, alongside other supervised methods [53–55], has shown significant success in addressing the missing wedge problem. However, these data-driven approaches face limitations: first, they require computationally intensive supervised pretraining; second, they rely on WBP reconstructions that already exhibit missing wedge artifacts; and third, supervised learning techniques can be prone to generating fictitious densities and inaccurately positioning structures within the reconstruction [56, 57]. Supervised learning applications in tomography, including those using the U-Net architecture [52], have been documented to result in such feature hallucinations and misplacements [58, 59]. These problems are exacerbated when the training data is limited; thus, in cryoET, it could lead to misinterpreting rare events [60, 61] that are scantly represented in large datasets.

As an alternative, we explore a novel unsupervised learning strategy [62–64] that bypasses the limitations associated with supervised learning [56–59] and the reliance on artifact-prone WBP reconstructions. Our approach starts with a randomly initialized network, optimizing its weights so that the generated image agrees with the experimentally captured projections, thus avoiding the need for pretraining on compromised WBP reconstructions with missing-wedge-induced artifacts. We employ coordinate networks (CNs) [65] to reconstruct this unsupervised representation of the tomogram. The CN determines 3D voxel values in the reconstruction volume by relating them to the corresponding 2D pixels in the projection images. Unlike conventional kernel-based methods such as CNNs, CNs offer a continuous representation by mapping coordinates to their corresponding values through a network-embedded continuous function—this allows CNs to capture image details without being constrained by a fixed grid resolution. Given their growing application in computationally intensive tasks in computer graphics [66, 67] and demonstrated potential in various biomedical imaging applications [68–73], CNs present a powerful solution for accurately representing extensive cryoET volumes, addressing the challenges of high computational costs and fixed-resolution limitations associated with CNNs.

Our study reveals that unsupervised learning with CNs can enhance shape fidelity and diminish the impact of the missing wedge on *in silico* data compared to traditional and CNN-based methods. Furthermore, bypassing the pretraining step allows CNs to produce reconstructions between three to over twenty times more rapidly than pretrained CNN methods. To rigorously assess image quality, we employed various voxel-based metrics and introduced a novel directional Fourier Shell Correlation (FSC) metric. This new metric is tailored to specifically quantify the restoration of information within the regions affected by the missing wedge.

While our findings highlight certain advantages of unsupervised learning for cryoET reconstruction, they are preliminary and not intended to establish superiority over other methods. Instead, we compare traditional WBP and Fourier inversion reconstructions against supervised and unsupervised machine learning frameworks, shedding light on their respective benefits and limitations within the broader context of structural biology and molecular imaging. Through this comparison, we aim to contribute valuable insights into the ongoing discourse of cryoET reconstruction techniques.

This manuscript is organized as follows: Section 2 describes the cryoET forward model (Figure 1), our reconstruction algorithm (Figure 2), data, experimental setup, and evaluation methods. Qualitative and quantitative results (Figures 3, 4, 5) in Section 3 are followed by a discussion in Section 4. Additional results can be found in the Appendix.

**Figure 1.**
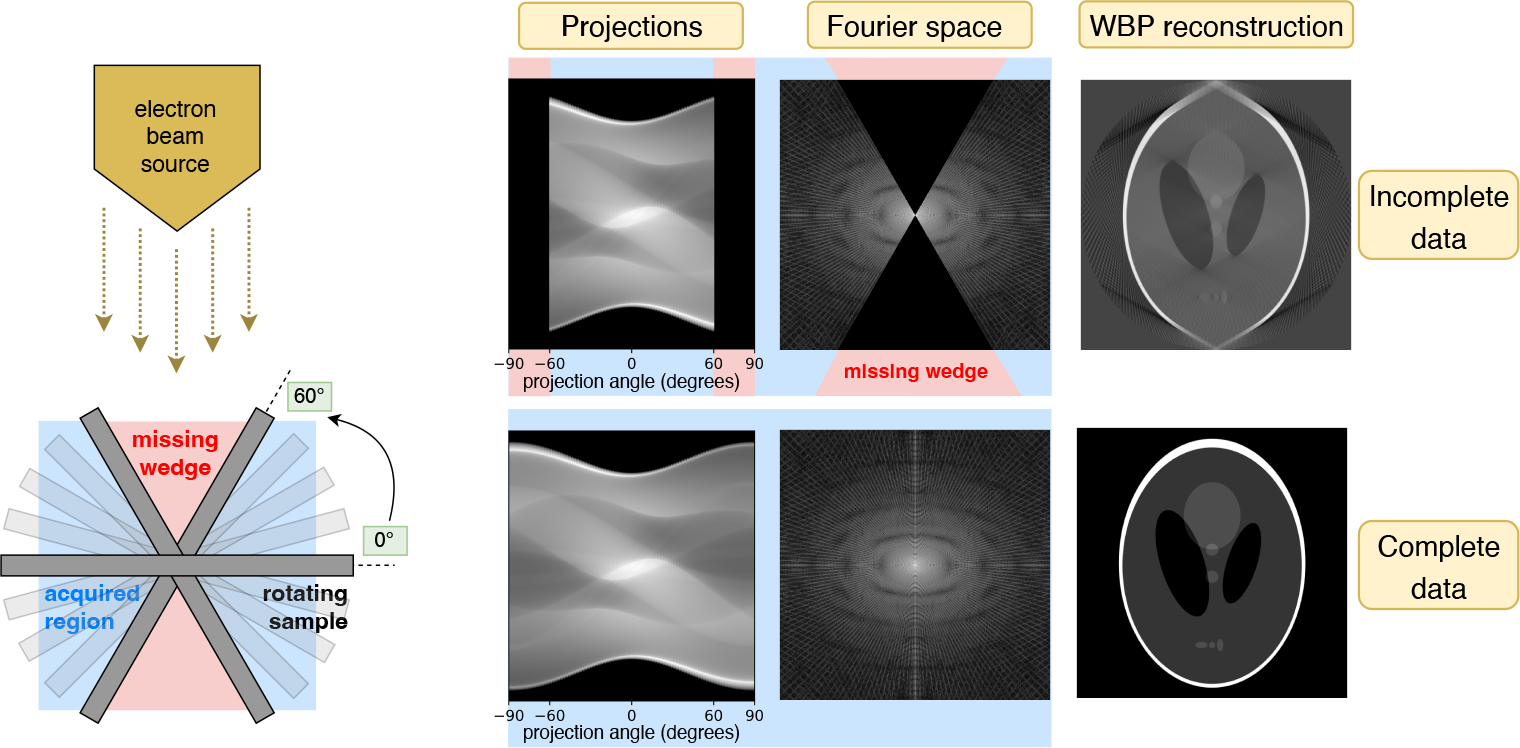
**<u>Left:</u>** CryoET acquires tomograms by collecting a tilt series of 2D projection images of vitrified biological samples over a limited angular range. **<u>Right:</u>** Simulations of the widely used Shepp-Logan phantom model show that restricted projection angles result in a missing wedge in Fourier space, leading to distortions in the reconstructed image (top row). The objective of this study is to reconstruct the uncollected data in this region, effectively completing the wedge (bottom row).

**Figure 2.**
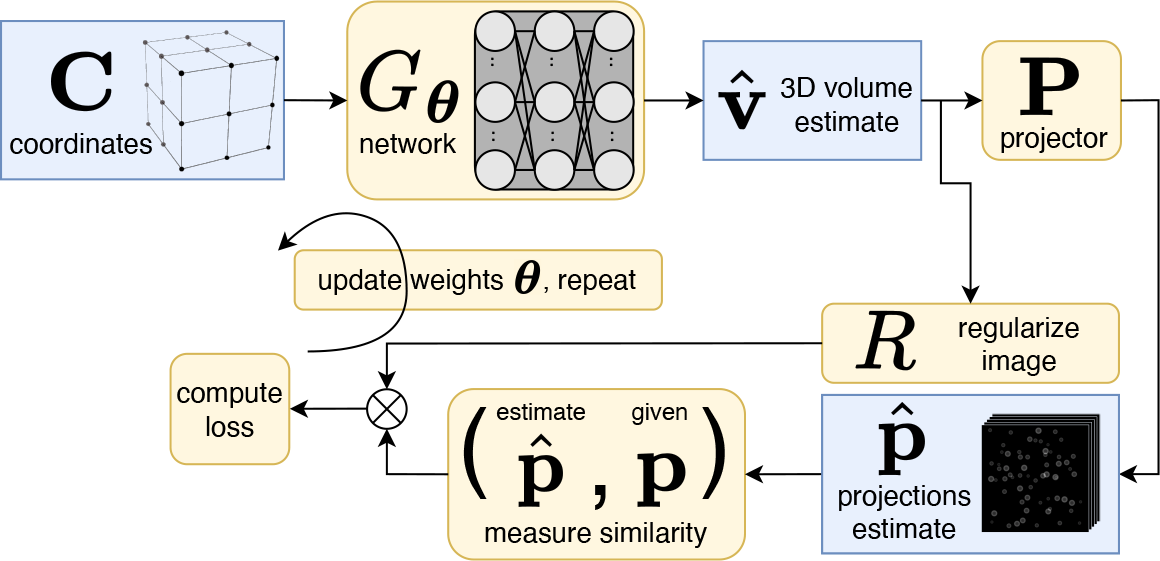
Unsupervised reconstruction using a coordinate network (CN) to create a 3D volume estimate, 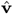. This diagram depicts one iteration of training process, where network weights ***θ*** are updated by constraining the estimated projections 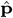 to match the given projections, **p**. After the training process, 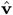 closely matches the ground-truth tomogram.

**Figure 3.**
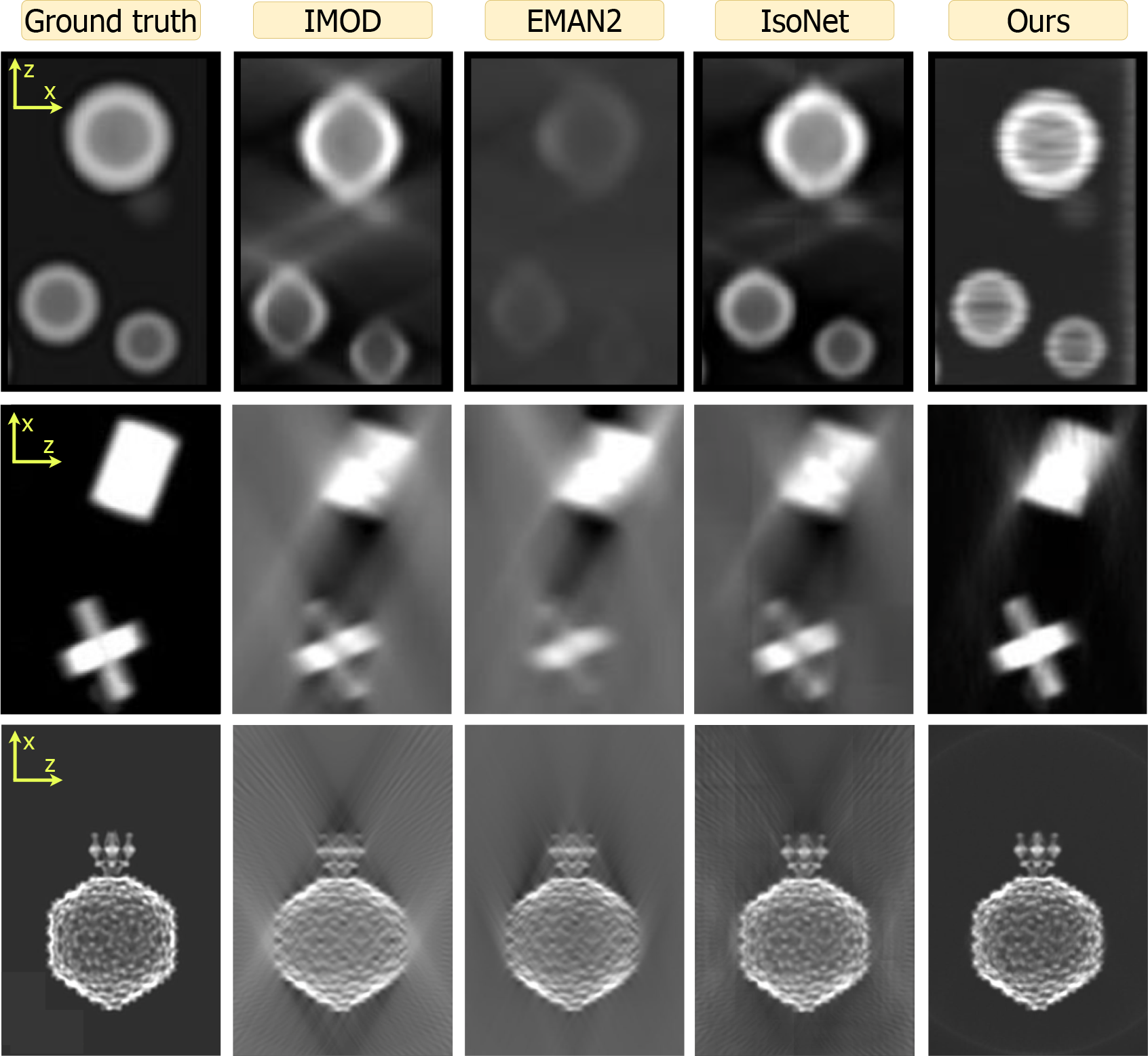
Qualitative review of the spheres (top), shapes (middle), and P22 (bottom) datasets across all methods (columns). Images are zoom insets of projections in the xz-plane. Both IMOD and EMAN2 suffer from back-projection artifacts. Compared to IsoNet, our method better resolves these artifacts and produces higher shape fidelity. Both IsoNet (tile pattern) and ours (horizontal streaks) can produce artifacts due to computational constraints, as discussed in Section 4. Figures A1, A2, A3 contain all projection directions and a larger field of view.

**Figure 4.**
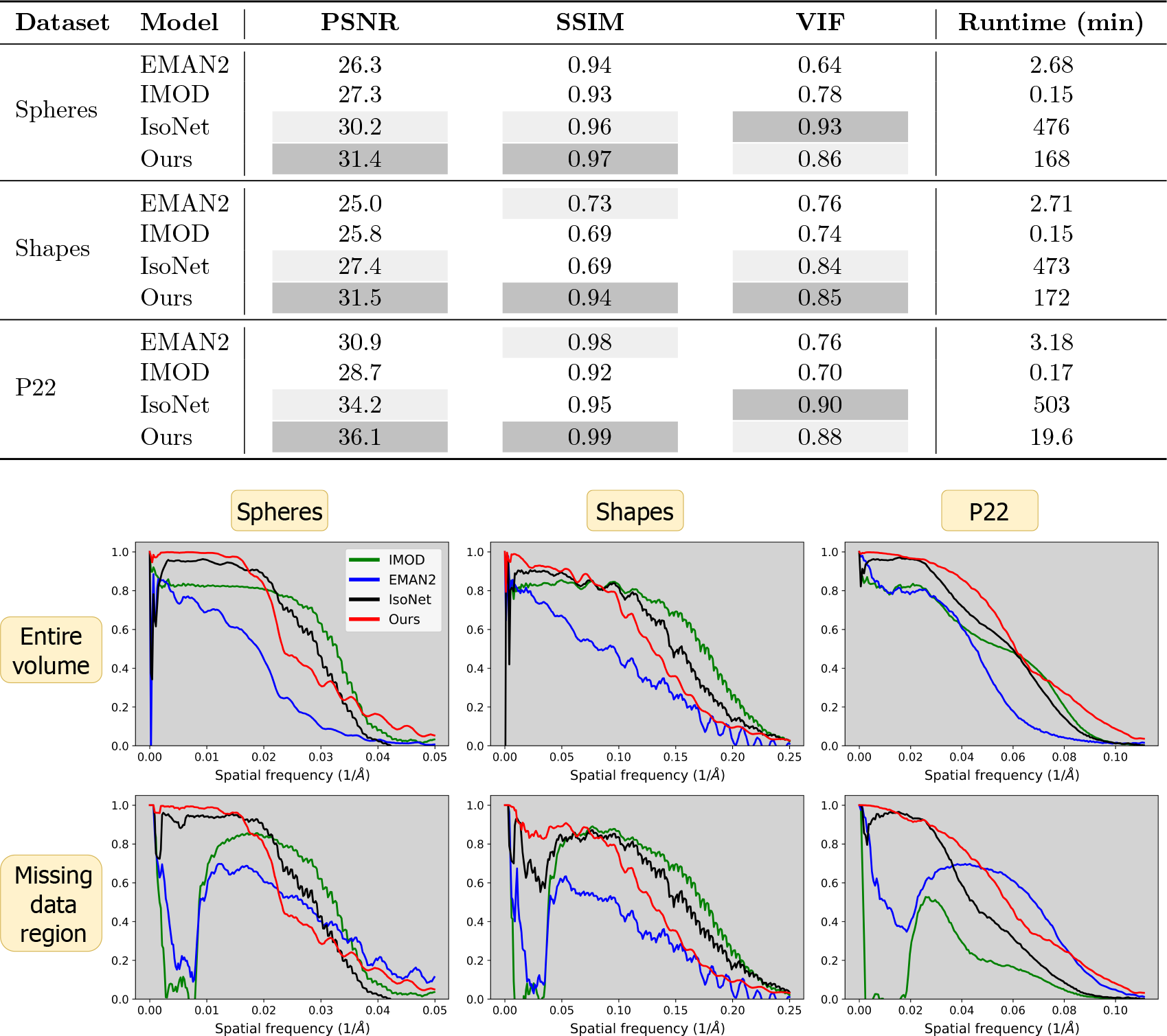
Image quality metrics across all datasets and models. **<u>Top:</u>** Voxel-based metrics and algorithm runtime. Ours (CN) generally performs best except for the VIF metric, which measures high frequencies. Due to pretraining, IsoNet is the most computationally intensive. **<u>Bottom:</u>** Fourier shell correlation (FSC) curves for each dataset (columns) across the entire volume (top row) and missing data region (bottom row). Ours typically performs best at lower frequencies and is surpassed at higher frequencies. Only ours and IsoNet provide substantive information at lower frequencies.

**Figure 5.**
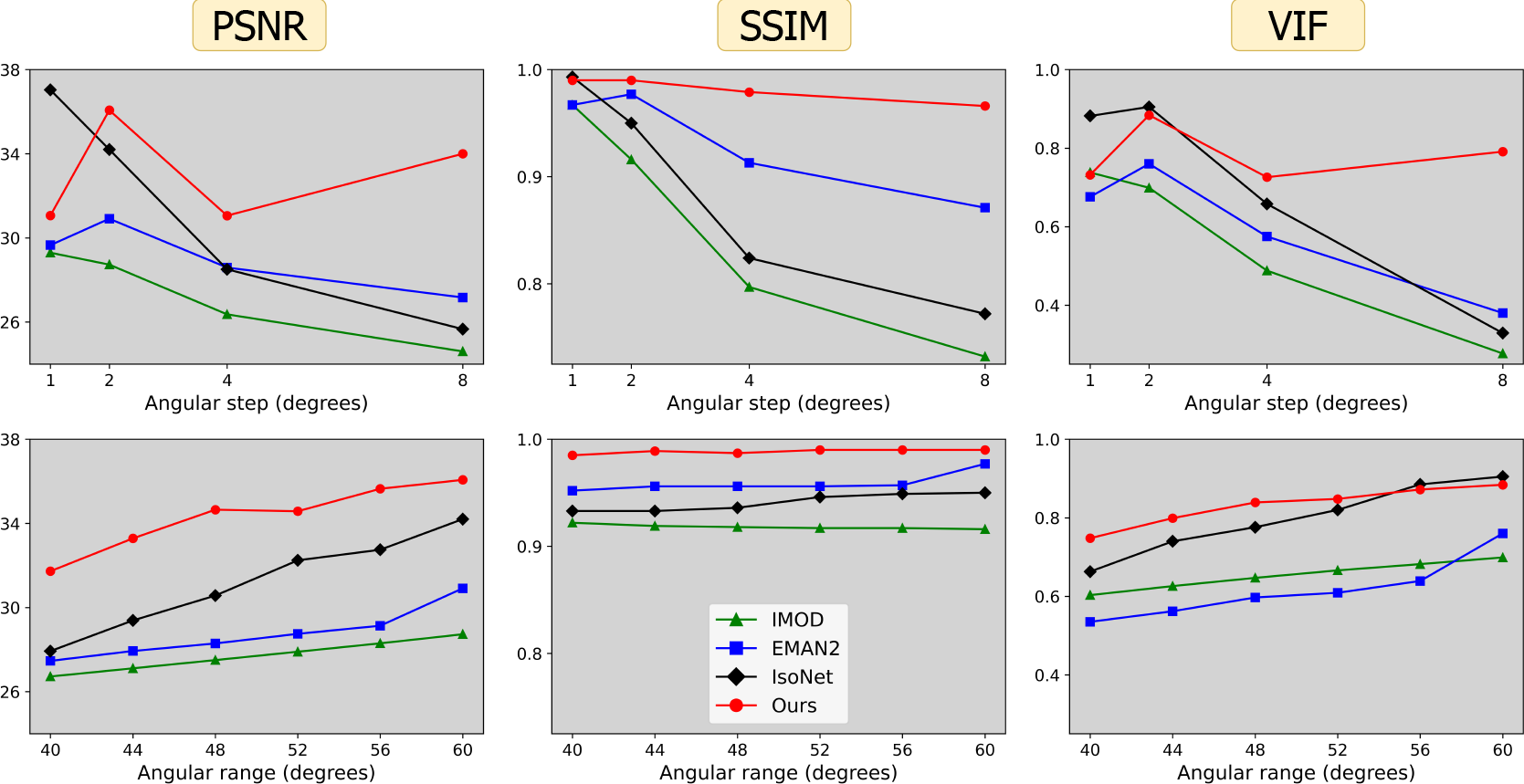
Voxel-based metrics (columns) with varying acquisition parameters: angular step, *i*.*e*. number of degrees between tilts *α* (top row) and angular range *β*, corresponding to projections over the range of [*−β, β*] degrees (bottom row). Default acquisition parameters are *α* = 2 and *β* = 60 degrees.

## 2 Materials and Methods

### 2.1 Forward Model

Let **v***\* ∈* ℝ^*x×y×z*^ be the true image volume (tomogram) we wish to reconstruct given access to projections **p** *∈* ℝ^*l×x×y*^, *i*.*e*. **p** = **Pv**^*\**^. Here **P** denotes the projection operator, which projects electrons through the volume **v***\** in a parallel beam at *l* different tilt angles. This process provides **p**, a tilt series of *l* projection images, each size *x × y*.

### 2.2 Reconstruction Algorithm

We employ a CN *G*_***θ***_ : ℝ^3^ *→* ℝ with trainable parameters ***θ*** that maps an individual 3D coordinate **c** *∈* ℝ^3^ in the reconstruction volume to a pixel value at the corresponding locations in the 2D projection images. Evaluating this network over the entire set of coordinates **C** = *{***c**_*q*_*}*^*x×y×z*^ produces a reconstruction volume *G*_***θ***_(**C**) *∈* ℝ^*x×y×z*^.

Our goal is to find a set of parameters for the CN such that its reprojections—the projector applied to the network output, *i*.*e*. **P***G*_***θ***_(**C**)—matches the experimentally given projections **p**. Hence we randomly initialize parameters ***θ*** and solve the following:

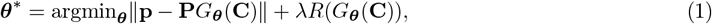

where *R* is a regularization term applied to the estimated image with strength *λ*. Because the projection operator **P** is differentiable, we can use gradient-based backpropagation to solve this equation. Figure 2 depicts this training process. The resulting network then produces 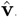, an estimate of the image volume we wish to reconstruct, *i*.*e*. 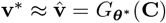.

Our methodology distinguishes itself by employing an unsupervised approach, thereby obviating the necessity for pretraining. Contrary to pretrained strategies that depend on supervising with data augmentation strategies—such as using subtomograms that replicate the effects of the missing wedge—our technique refines network parameters by directly using the experimental projection images. This direct optimization method effectively circumvents the common pitfalls of supervised learning, including the propensity for some types of artifact generation and structural inaccuracies [56–59]. Indeed, our approach leverages more dependable data—the experimental projection images themselves—avoiding the artifact-laden WBP reconstructions commonly utilized as a starting point for supervised methods. By ensuring a closer agreement between the reconstructions’ reprojections and the original experimental projections, our method inherently reduces the likelihood of introducing hallucinated errors or artifacts.

### 2.3 Data

To compare our approach with other reconstruction methods, we carried out *in silico* experiments for which ground truth is known, enabling precise evaluation via quantitative, reference-based metrics. Tomograms were created with image processing tools available in EMAN2 [74], and their corresponding tilt series were generated by projecting through the volumes every 2° across the range of -60° to +60° using EMAN2’s standard projector. See below for descriptions of each dataset (*x × y × z*):

#### Spheres

(1024 *×* 1024 *×* 256): a collection of binarized hollow spheres of constant density and variable sizes (16 to 64 pixels in diameter). Compare to *x* and *y*, the smaller dimension in *z* renders a slab-shaped volume, geometrically mimicking a distribution of discrete objects in a thin layer of ice.

#### Mixed shapes

(1024 *×* 1024 *×* 256): varied geometric shapes with heterogeneous structures. These binarized shapes include full spheres, ellipsoids, pyramids, cubes, rectangular prisms, circular discs, as well as 4- and 6-pointed 3D crosses. Similar to spheres, the slab shape of this tomogram mimics the geometry of a thin layer of ice.

#### P22

(360 *×* 360 *×* 360): single P22 phage particles. The P22 capsid displays icosahedral symmetry, while the virion tail exhibits pseudo-six fold symmetry. This map, accessed via the electron microscopy data bank (EMDB, accession number EMD-9008) [75], is sampled at 4.5 Å/pixel. We clipped the volume to a 360 *×* 360 *×* 360 box size and threshold filtered it to eliminate negative densities.

#### Ubiquitin

(64 *×* 64 *×* 64): the regulatory protein ubiquitin. We created this simulation using a map generated from an atomic model downloaded from the protein data bank (PDB)(PDB ID: 1UBQ) [76].

For final processing steps, each simulated tomogram (except for ubiquitin) was low-pass filtered to either render shape surfaces smooth (spheres, shapes) or dampen high-resolution features that would not be present in a raw cryoET tomogram without averaging (P22, ubiquitin). Finally, each tilt series of projection images was normalized to be within the range of [0, 1] to standardize input values provided to the network.

### 2.4 Experimental Setup

To find a set of weights ***θ***^*\**^ that minimize Equation 1, we construct a fully-connected coordinate network architecture in PyTorch [77]. This network has four hidden layers each with 256 features, positional encoding [66], and sinusoidal activation functions [65]. For simplicity, we employ this same architecture on all datasets and maintain a consistent ratio of network parameters to measurements (roughly 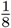). This consistency is achieved by dividing the tilt series **p** *∈* ℝ^*l×x×y*^ into length-*j* subslices along the *y*-axis, *i*.*e*. **p**_sub_ *∈* ℝ^*l×x×j*^; in image space, this corresponds to a subvolume 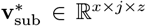. Subsequently we fit a separate set of network parameters to represent each subvolume. For example, given the aforementioned network with 4 *\** 256^2^ = 262, 144 parameters and a tomogram of size 1024 *×* 1024 *×* 256, obtaining a 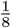 ratio would yield 128 networks, each fitting measurements for a subvolume of size 1024 *×* 8 *×* 256. We then stitch these individual subvolumes together along the *y*-axis to obtain the final reconstructed volume.

Given sufficient memory, a sparser representation, or a smaller volume, one could reconstruct the entire volume with one single network as we demonstrate in Figure A4. However, we find our subvolume approach has several advantages: (1) it ensures the network has sufficient representational capacity for any tomogram, (2) it uses a small amount of memory–––roughly 2-4 GB on our NVIDIA Quadro RTX 8000— making this method feasible on smaller GPUs, and (3) it enables us to leverage learned initializations [68], *i*.*e*. after fitting a network to one subvolume, those same parameters are used to initialize the network for the adjacent subvolume. This learned initialization strategy improves reconstruction quality by enhancing consistency along the *y*-direction and also reducing the number of gradient step iterations required for the network to fit the subvolume. As such, we use 2,000 iterations to fit the first subvolume and 400 iterations for all adjacent subvolumes which leverage learned initializations. For the first subvolume, we use an initial learning rate of 1 *×* 10^*−*3^ decayed logarithmically to 1 *×* 10^*−*4^; for all adjacent subvolumes, we use an initial learning rate of 1 *×* 10^*−*4^ decayed logarithmically to 1 *×* 10^*−*5^. By default, *λ* = 0 in Equation 1.

### 2.5 Baselines

We compare our algorithm to three well-established baselines previously introduced in Section 1: (1) WBP [41] reconstructions generated via IMOD [42], (2) Fourier inversion reconstructions generated using EMAN2 [78], and (3) reconstructions with missing wedge restoration by IsoNet [79], a supervised deep learning approach leveraging CNNs to fill in the missing wedge of the IMOD reconstruction provided as input. We choose IsoNet because it is widely regarded as the leading method for missing wedge compensation, outperforming approaches such as ICON [51] or MBIR [49]. We use default parameters for each reconstruction method.

### 2.6 Image Evaluation

To provide a comprehensive evaluation of signal preservation across frequencies in these reconstructions, we employ the Fourier shell correlation (FSC) metric, common in structural biology, as well as voxel-based metrics from the computational imaging literature. All metrics are reference-based, comparing the reconstructed image 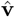 against a ground truth reference image **v***\**. For each metric, higher values indicate superior quality.

#### 2.6.1 Directional Fourier shell correlation (FSC)

FSC measures the similarity between the Fourier coefficients of a reconstructed image volume and those of its ground truth reference. This measurement is performed by selecting a specific radius in Fourier space and identifying points within a half-unit distance from the sphere’s surface corresponding to that radius. These identified points contribute to the calculation of a normalized Pearson correlation coefficient, which is computed without subtracting the mean. The process involves incrementally adjusting the Fourier radius from one unit up to the Nyquist frequency, calculating the Pearson correlation at each step to assess the correlation across different spatial frequencies.

We first apply the conventional FSC metric across the entirety of the reconstructed volumes. Additionally, we introduce a directional FSC variant designed specifically for assessing the missing data regions. This innovative approach aims to directly highlight the effectiveness of a reconstruction algorithm in compensating for the missing wedge. We characterize the “present data” region as the areas within a half-unit distance from any plane defined by the direction of the *l* acquired projections, effectively defining a slab for each tilt angle; the “missing data” regions correspond to the complement of the present data. While alternative methods for interpolating data within this geometric framework are possible, the FSC typically exhibits a smooth profile across these calculations. Consequently, we anticipate similar results with varying interpolation strategies.

#### 2.6.2 Voxel-based metrics

This subsection outlines three metrics widely recognized in imaging research, each assessing distinct aspects of image quality.

##### Peak Signal-to-Noise Ratio (PSNR)

quantifies the ratio between the maximum possible power of a signal and the power of corrupting noise that affects its representation. Measured in decibels, PSNR is derived from the mean-squared error (MSE) between a reconstructed image and its ground truth. The formula for PSNR is:

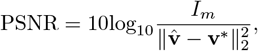

where *I*_*m*_ represents the maximum possible pixel value (*e*.*g*., 255 in an 8-bit grayscale image) and the denominator corresponds to MSE. This metric is particularly suited to measuring similarity of low-frequency image components [80].

##### Structural Similarity Index (SSIM)

[81] measures perceived image quality by evaluating aspects like structure (texture and pattern consistency), luminance (brightness levels), and contrast (voxel variance). SSIM values range from -1 to 1, with 1 signifying perfect similarity, *i*.*e*. 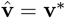. Compared to PSNR, SSIM offers a more nuanced evaluation of image quality, closely aligning with human visual perception [82].

##### Visual Information Fidelity (VIF)

[83] quantifies how well the reconstructed image captures natural scene statistics corresponding to the human visual system (HVS). It measures mutual information (MI) between the input and outputs of both the reconstructed and reference volumes, **v** and **v***\**. Hence the formula for VIF can be written as:

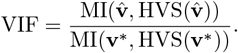

Possible values of this metric can be between 0 and 1 (blurry 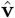), 1 (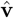 = **v**\*), or greater than 1 (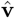 provides contrast enhancement of **v***\** without adding noise). This metric best captures similarity between higher frequency components of an image. In contexts such as magnetic resonance imaging, VIF has demonstrated alignment with radiologist preferences [84].

## 3 Results

Figure 3 shows *xz*-plane reprojections of small regions of the spheres and shapes datasets as well as the full span of a P22 volume. All orthogonal reprojections are shown in Figures A1, A2, and A3 with the *xz*-projection trimmed in size for display purposes. The missing wedge artifact due to anisotropic resolution manifests as elongation streaks in the *xz*-plane and blurriness along *z* in the *yz*-projections. Across methods, IMOD and EMAN2 reconstructions exhibit these artifacts most strongly. Such artifacts are substantially reduced by IsoNet and the least pronounced in reconstructions using our approach.

We now consider the accuracy of shape representation. For the spheres dataset, both IMOD and EMAN2 inaccurately render ellipsoidal contours elongated along the *z*-axis. For the P22 dataset, these methods fail to achieve the expected sharpness along the edges of the particle in *z*. Across all datasets, IsoNet reduces these distortions considerably. Meanwhile, our CN reconstruction preserves shape fidelity closest to ground truth, reducing distortions more than IsoNet.

Figure 3 exhibits high-frequency reconstruction artifacts for IsoNet in the form of tiling (spheres, shapes) and for our method in the form of high frequency streaks (spheres). In spite of this, both IsoNet and our method preserve overall shape fidelity at low and intermediate frequencies, which are most relevant in cryoET outside of high-resolution STA applications. We provide a thorough discussion of these artifacts in Section 4.

To quantitatively evaluate reconstruction quality, we utilized several voxel-based metrics alongside the FSC (Figure 4). Our technique outperforms others in terms of PSNR, which assesses performance at lower frequencies, and SSIM, which evaluates structural integrity. However, IsoNet outperforms our method in VIF, indicative of higher frequency accuracy, in two out of the three datasets examined. This observation is consistent with FSC analysis, which demonstrates superior performance of our method at lower frequencies while trailing at higher frequencies. FSC curves for the missing data regions highlight the proficiency of both our method and IsoNet in compensating for the lack of information in the missing wedge compared to IMOD and EMAN2.

To further test robustness of different methods, we generated alternate tilt series from one of the P22 tomograms by varying two critical parameters: angular step (*α*) and angular range (*β*). CryoET data collection often uses smaller tilt steps of 1-2° for large, continuous specimens such as cells; however, tilt steps of 3-5° or larger can be used for sparse specimens such as macromolecules in solution destined to undergo STA [33]. Similarly, data collection ranges can be smaller than [-60°, +60°] for high-resolution STA, as high-tilt images may be too noisy and damaged by cumulative radiation dose.Figure 5 shows the impact of variations in *α* and *β* on the performance of each reconstruction method. Our findings affirm the adaptability of our method to different acquisition parameters, underscoring the potential utility of our approach in unique applications requiring varied or non-standard data collection parameters.

## 4 Discussion

Our CN reconstruction method exhibits superior performance in low-frequency ranges as depicted in Figure 4, a trait consistent with observations in unsupervised learning methods noted for their low-frequency spectral bias [62–64, 85–87]. This may be advantageous for cryoET tasks that require shape integrity such as feature segmentation and particle picking—which can be challenging, slow, and inconsistent in typical cryoET tomograms due to missing-wedge-induced resolution anisotropy [88, 89]. At higher frequencies, our method performs worse than IMOD and IsoNet as deemed by the VIF metric and FSC curves in Figure 4. This suggests that our method may not benefit high-resolution applications, such as completing the missing wedge in single-particle cryoEM analyses of macromolecular complexes which exhibit preferred orientation; for this problem, various experimental [90] and algorithmic [91–93] approaches have been proposed. The fact that different methods perform better in different frequency ranges prompts considering an ensemble approach for future advancements, potentially integrating our unsupervised model’s strengths in lower frequencies with IsoNet’s proficiency in higher frequency details, to provide a more uniformly high-quality reconstruction.

The varied performance across frequency ranges also underscores the critical role of tailored evaluation metrics, such as our innovative use of directional FSC to evaluate information in the missing data region. This specific assessment underlines the advantage offered by neural network approaches like ours and IsoNet in compensating for the missing wedge. Given that no single reconstruction method performs best across all frequencies or samples, selecting varied test datasets and evaluation metrics is essential for a comprehensive assessment of reconstruction methods.

Our direct use of projection images bypasses the initial distortion introduced by WBP images that IsoNet uses for training. We hypothesize this promotes a higher fidelity to the original shapes within the tomograms, as exemplified by our spherical outcomes versus IsoNet’s ellipsoidal tendencies in Figure 3 and by our superior performance at low frequencies and in structural similarity 4. This also reveals a trade-off in our reconstruction approach, as demonstrated in Figure 5. Our method’s reliance on projections means that variations in the angular step (*α*) and range (*β*) directly alter the volume of information we process. In contrast, IsoNet’s dependency on WBP images means that, while changes in *α* and *β* affect the quality of its WBP input image, the amount of input data—essentially the image size —remain unchanged. Interestingly, when we decrease *α* to 1°, ostensibly increasing the available information, our method’s performance unexpectedly dips. This likely stemmed from our network’s capacity being held constant throughout these experiments, despite processing an increasing amount of projections with decreasing tilt step; i.e., the network capacity may have been exceeded going from 2° to 1°. This observation motivates our decision to dynamically adjust the network size in response to the dimensions of the tomogram and the number of projections available, as discussed in Section 2.4. This adaptability ensures that the network’s representational capacity—essentially the ratio of network parameters to tomogram size—remains consistent. Such consistency is beneficial for achieving uniform quality across reconstructions of varying sizes and complexities. Additionally, this feature facilitates a scalable runtime, proportional to the tomogram’s size. For instance, when transitioning from the larger spheres dataset to the smaller P22 dataset, we observed an 89% decrease in runtime (Figure 4), closely mirroring an 83% reduction in tomogram volume. In contrast, IsoNet exhibits a relatively uniform runtime across these datasets, highlighting a fundamental difference in our approaches.

Our method’s capacity to adjust network size allows for customized reconstruction scale based on the hardware available: smaller subvolumes along the *y*-axis on more modest systems, and larger subvolumes on more powerful setups. In principle, dividing the tomogram into adjacent subvolumes to meet memory constrains of the GPU also allows for parallelization to derive future speed gains. However, this advantage does not come without challenges. Specifically, it can lead to the emergence of streaking artifacts along the *y*-axis, a phenomenon observable in the spheres reconstruction depicted in Figure 3. The occurrence of such artifacts, however, is not inherent in our CN methodology, as demonstrated by our experiment using the small regulatory protein ubiquitin in Figure A4, which shows these artifacts vanish when a single network is employed to reconstruct the entire volume. Yet, this single-network approach is not currently feasible for larger datasets like our spheres and geometric shapes simulations due to memory constraints on our hardware (NVIDIA Quadro RTX 8000). Future hardware developments or parallelization could alleviate this issue. Separately, future algorithmic improvements could more efficiently represent large volumes, using strategies such as Gaussian splatting [94] or Gaussian mixture models, as demonstrated for SPA cryoEM [95, 96].

IsoNet also grapples with computational limitations inherent to 3D CNNs, resulting in tiling artifacts in reconstructions (Figure 3). These artifacts stem from the necessity of dividing the training set into manageable 3D subtomograms, each measuring 64 *×* 64 *×* 64 voxels by default. This methodological constraint underscores a shared challenge in cryoET reconstruction: balancing computational feasibility with the aim of computing artifact-free, high-fidelity reconstructions.

While our *in silico* results are promising, real-world application remains under development, presenting a frontier for future research. Real datasets introduce complexities such as noise and CTF, necessitating advanced unsupervised learning techniques for effective noise management and image enhancement [97, 98]. Indeed, the CTF is particularly challenging to correct for in cryoET datasets due to the defocus gradient present in cryoEM images of tilted specimens [99, 100], which worsens with increasing specimen thickness [101], tilt angle [102], and field of view such as at lower magnifications. CTF-induced artifacts can limit the resolution at which macromolecules and biological specimens in general can be visualized at, and artificially give an appearance of hollowness to solid objects depending on their shape, size, and the defocus amount. Moreover, IsoNet’s comprehensive image processing pipeline, which includes preprocessing steps before reconstruction, such as CTF deconvolution [103] and masking, underscores the challenges to making objective, holistic comparisons between reconstruction methods. Here, we focus on characterizing the performance of unsupervised machine learning methods and evaluating trade-offs with supervised ones.

Given the increasing popularity of cryoET [104] owing to its demonstrated applications in cellular structural characterization [105] and histopathological clinical diagnoses [32], we expect for artificial intelligence developments that enhance tomographic reconstruction to become an increasingly active and impactful field of research. Since cryoET can provide 3D views of individual macromolecules and complexes across a wide range of sizes, as well as their distributions within organelles, cells, and tissues, improving tomographic reconstruction quality may enable novel structure-based diagnostics at the molecular level and accelerate drug development by assessing the effects of molecular therapeutics on phenotype.

## 5 Acknowledgments

This research was funded by a grant from the Chan-Zuckerberg Institute entitled “Resolving Biostructures In-situ via Cryogenic Light and Electron Microscopy” in addition to NIH grants U01 AI136680 and U54 AI170855.

## A Appendix

**Figure A1.**
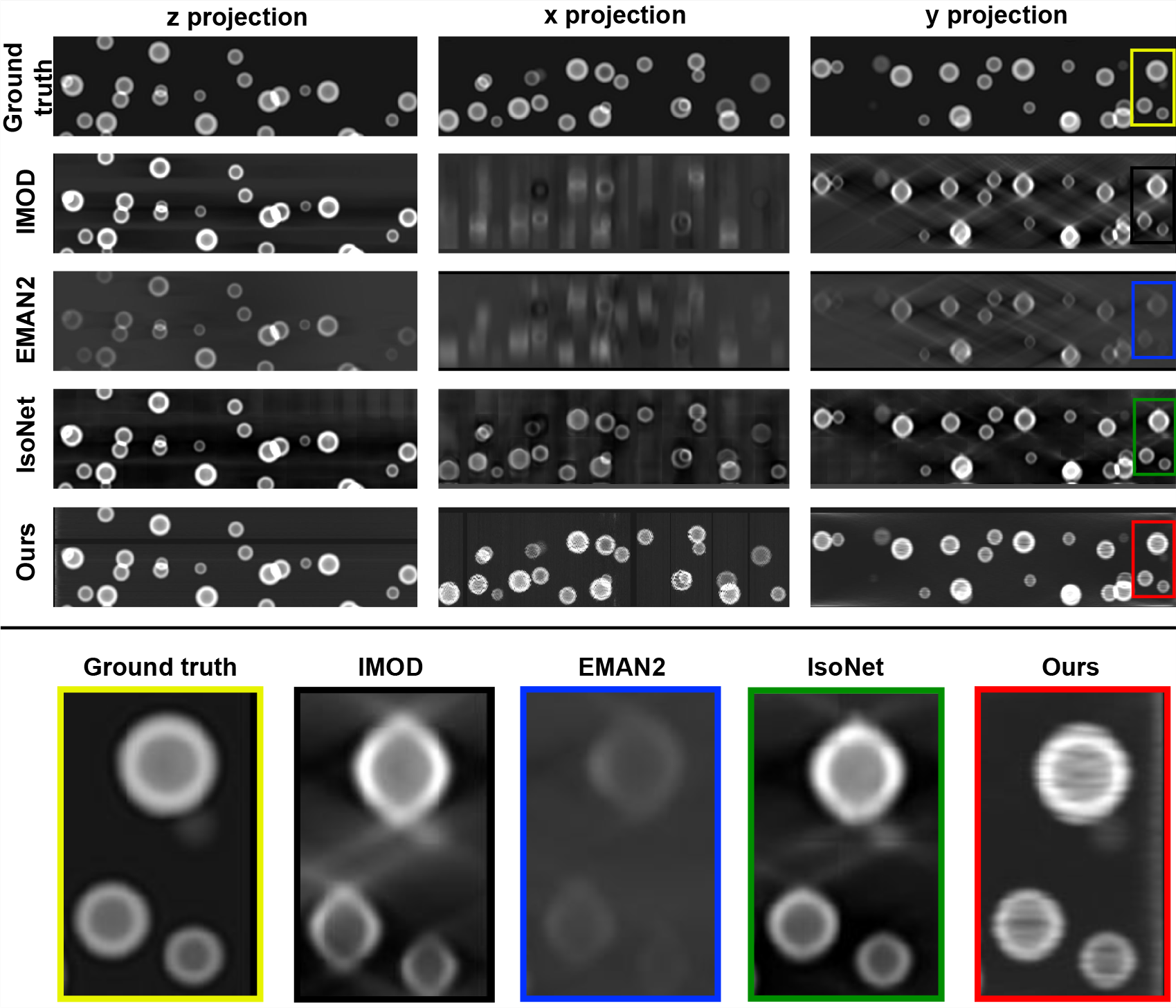
Qualitative review of the spheres dataset. **<u>Top:</u>** Projections in each direction (columns) for each method (rows). **<u>Bottom:</u>** Zoom inset of the xz-plane. IMOD and EMAN2 suffer from back-projection artifacts. Compared to IsoNet, ours better resolves these artifacts and also produces a more spherical shape. Reconstruction artifacts are present in both IsoNet (tile pattern) and our (horizontal streaks) reconstructions due to computational constraints, as discussed in Section 4.

**Figure A2.**
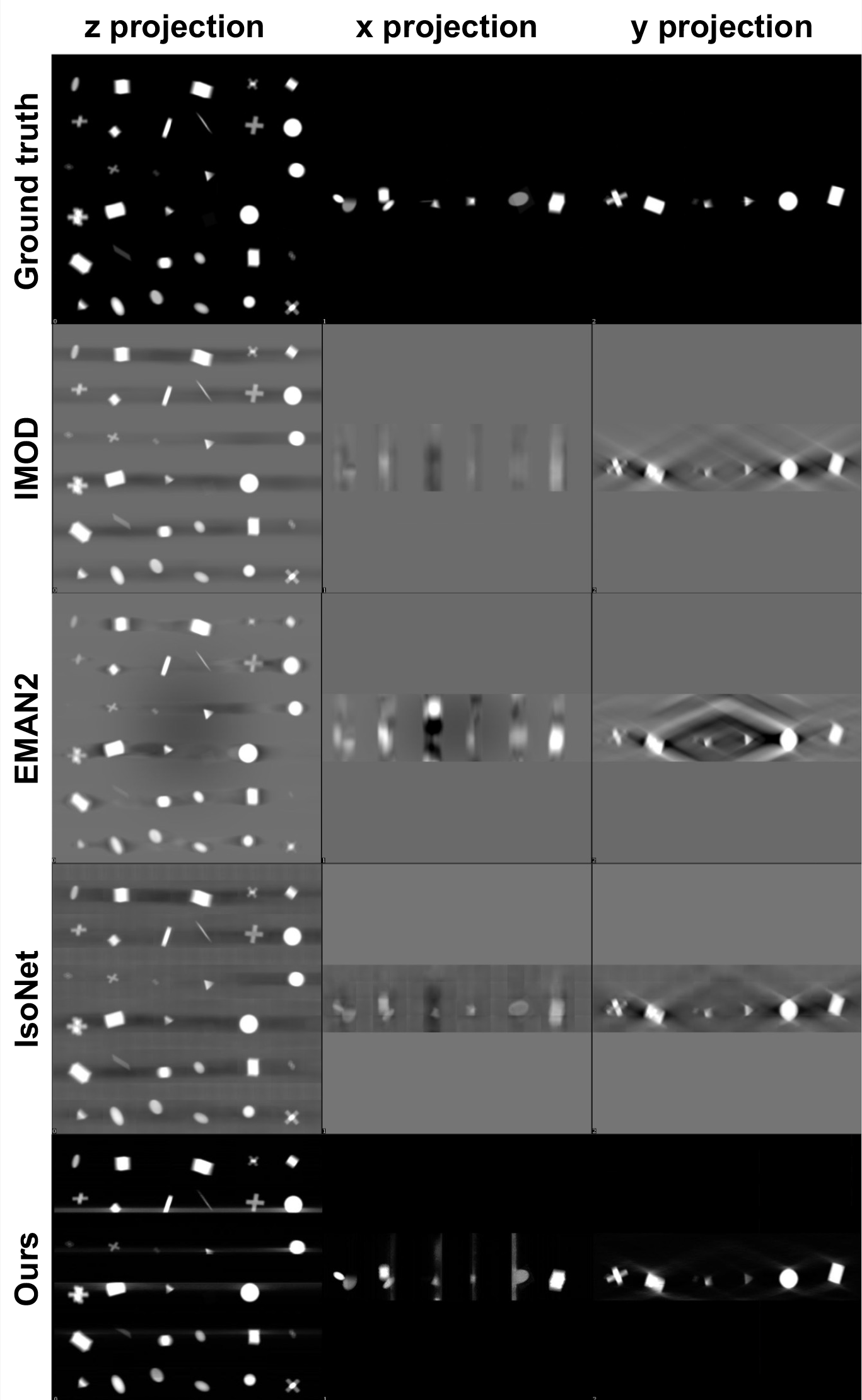
Qualitative review of the geometric shapes dataset: projections in each direction (columns) for each method (rows). Ours provides the best shape completion and mitigation of back-projection artifacts.

**Figure A3.**
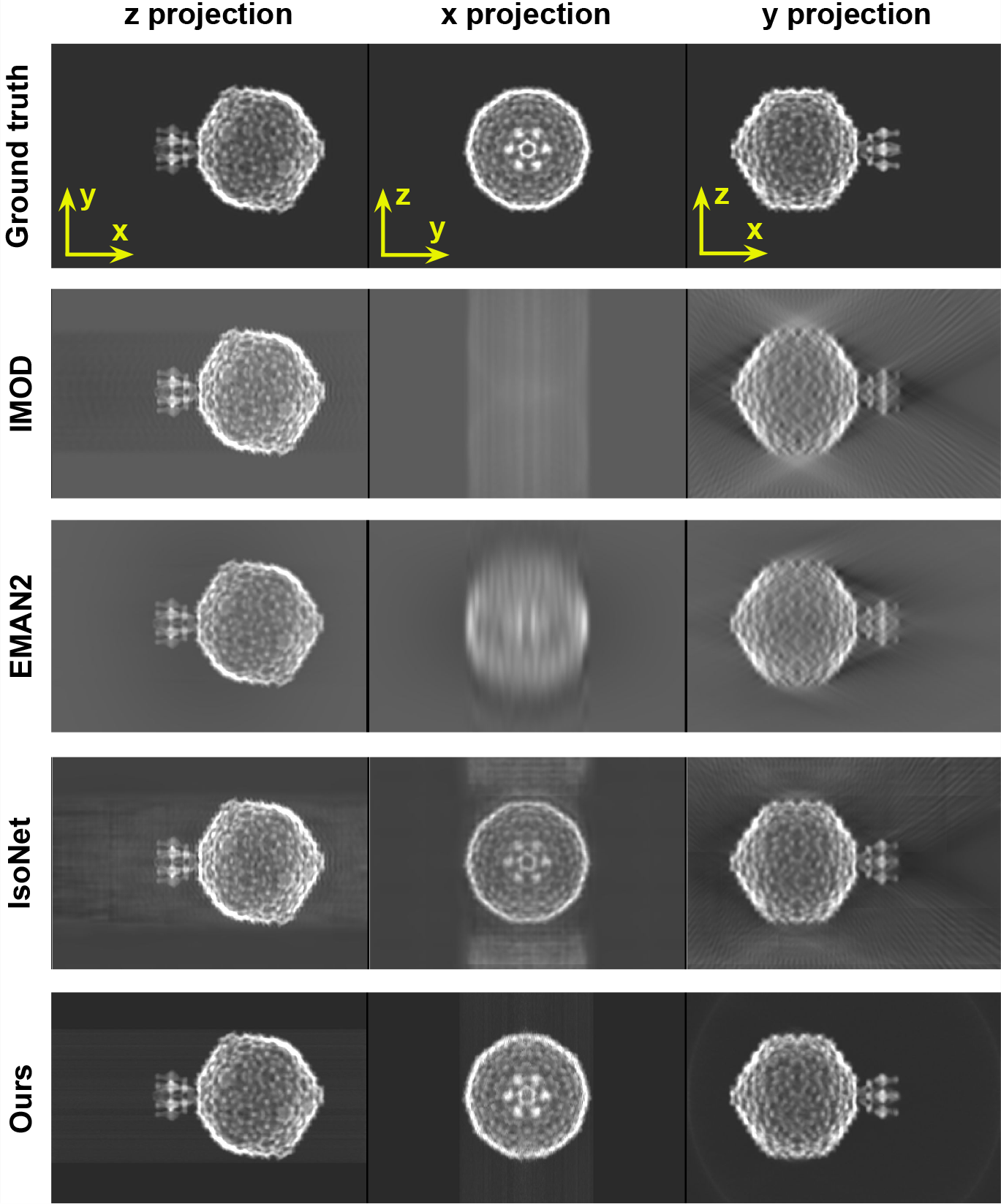
Qualitative review of the P22 dataset: projections in each direction (columns) for each method (rows). Ours provides the best shape completion and mitigation of back-projection artifacts.

**Figure A4.**
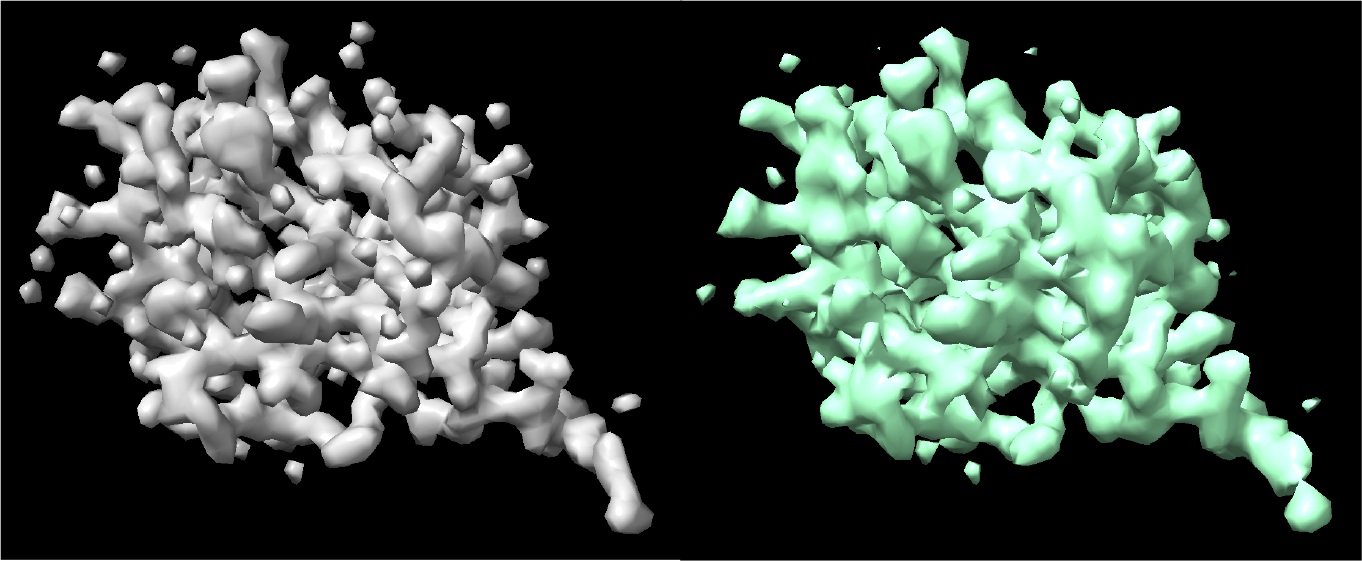
Qualitative review of the ubiquitin dataset comparing 3D volumes of the ground truth (left) vs. ours (right) to emphasize that streaky artifacts are absent when a single network is used to fit the tomogram, as discussed in Section 2.4. IsoNet also exhibits reconstruction artifacts due to computational constraints, as displayed in Figure 3. This motivates potential improvement for both methods and the development of hybrid approaches to more efficiently and accurately represent large cryoET volumes.

